# Synthetic design of farnesyl-electrostatic peptides for development of a protein kinase A membrane translocation switch

**DOI:** 10.1101/635839

**Authors:** Allen K. Kim, Helen D. Wu, Takanari Inoue

## Abstract

Molecular switches that respond to a biochemical stimulus in cells have proven utility as a foundation for developing molecular sensors and actuators that could be used to address important biological questions. Developing a molecular switch unfortunately remains difficult as it requires elaborate coordination of sensing and actuation mechanisms built into a single molecule. Here, we rationally designed a molecular switch that changes its subcellular localization in response to an intended stimulus such as an activator of protein kinase A (PKA). By arranging the sequence for Kemptide in tandem, we designed a farnesylated peptide whose localization can dramatically change upon phosphorylation by PKA. After testing a different valence number of Kemptide as well as modulating the linker sequence connecting them, we identified an efficient peptide switch that exhibited dynamic translocation between plasma membranes and internal endomembranes in a PKA activity dependent manner. Due to the modular design and small size, our PKA switch can have versatile utility in future studies as a platform for visualizing and perturbing signal transduction pathways, as well as for performing synthetic operations in cells.

## Introduction

Synthetic switch mechanisms are essential in developing molecular sensors and actuators with which scientists can probe biological phenomena by visualizing and perturbing underlying signal processes in living cells. Many of these switches rely on shifts in protein localization in cells that can be induced by post-translational modifications, such as phosphorylation at localization peptide sequences. For example, there are switches that exhibit translocation between cytosol and nucleus in response to phosphorylation by a target kinase^1–3^. In contrast, switches that shift between subcellular locations other than cytosol and nucleus are rare. In nature, we find cases where proteins change localization between two membrane systems. In the case of MARCKS and Src kinase, phosphorylation by protein kinase C (PKC) and both PKC and protein kinase A (PKA), respectively, could alter their subcellular localization (plasma membrane vs. cytosol)^4,5^. Developing a membrane switch would be useful in expanding the palette of molecular probes.

PKA is a cAMP-dependent kinase that plays a fundamental role in multiple cell functions including regulation of sugar metabolism, muscle contraction, heart rate, and blood pressure. There have been multiple PKA biosensors developed based on various switch mechanisms^6^. However, a molecular switch that shifts its localization within the membrane systems based on the PKA activity has yet to be developed.

To generate a synthetic molecular switch that translates the phosphorylation by PKA into its membrane localization, we looked into the regulatory mechanism of K-Ras4b. The localization sequence of K-Ras4b is found in the C-terminus where the combination of polybasic residues and the CAAX motif (where C is cysteine, A is aliphatic residue, and X is any residue) targets the protein to the plasma membrane. The CAAX motif serves as a signal for enzymatic processing (involving farnesyltransferase, Ras-converting enzyme I, and isoprenylcysteine carboxyl methyltransferase) that leads to the attachment of a farnesyl group^7–9^. As farnesylation alone is not sufficient for plasma membrane targeting, many plasma-membrane-targeted farnesylated proteins including K-Ras4b also require the presence of a cluster of positively charged residues near the CAAX motif ^10–15^. Phosphorylation of serine-181 by Protein Kinase C (PKC) leads to a disruption of K-Ras4b’s localization at the plasma membrane^16,17^. The electrostatic interaction between the positively charged residues in the protein and the negatively charged lipids in the plasma membrane is responsible for biasing the trafficking of K-Ras4b to the plasma membrane^18^. Phosphorylation in this region removes this bias by disrupting the favorable electrostatic interaction, a mechanism known as the farnesyl-electrostatic switch (FES). Later studies indicated that K-Ras4b uses more complicated mechanisms than a simple electrostatic switching for versatile membrane binding^13^.

While PKC requires a cluster of positively charged residues for substrate recognition^19^, a PKA recognition motif within PKA substrate sequences consists of two sequential arginines^20,21^. By aligning the PKA substrate sequence in tandem with conjugation to a farnesylation sequence, we designed an amino acid sequence that functions as a PKA phosphorylation switch that shuttles between the plasma membrane and the endomembrane depending on the PKA activity.

## Results

Inspired by the farnesyl-electrostatic switch (FES) mechanism of K-Ras4B, we aimed to develop a synthetic molecular switch for PKA activity. FES can be distilled into the following components (Fig. 1A): 1) series of positively charged-residues that will electrostatically interact with negatively charged lipids in the plasma membrane, 2) phosphorylation sites that will change positive charge content and/or orientation to regulate signaling, and 3) CAAX motif that will serve as a recognition sequence for farnesylation to target the membrane system such as plasma membrane and internal endomembranes. In our design of the FES that responds to PKA, we employed a short PKA substrate sequence, Kemptide (LRRASLG), to serve as a source of phosphoserine^20^. We flanked a single lysine with the two Kemptide sequences, and a single leucine was removed from the second Kemptide sequence to maintain a consecutive sequence of positively charged residues. We then fused a farnesylation substrate recognition sequence (TKCVIM), previously demonstrated to be sufficient for farnesyltransferase processing^22,23^. This resulted in a peptide design (Fig. 1B). The peptide was termed FES-PKA for farnesyl-electrostatic switch regulated by PKA activity. FES-PKA was expressed as an EYFP fusion in HeLa cells and was enriched at the plasma membrane along with localization at endomembranes. Upon activation of PKA with a cocktail of 50 μM forskolin (FSK) and 100 μM 3-isobutyl-1-methylxanthine (IBMX), the fluorescent signal redistributed to favor the endomembranes (Fig. 1C, Supplementary Video 1). To quantify this signal redistribution, we measured fluorescence intensity of the endomembranes which were independently visualized by an endomembrane marker, mCherry-Cb5. The fluorescence intensity of FES-PKA at endomembranes at time 0 (F0) was then subtracted from the intensity values at each time point (ΔF). We then plotted ΔF/F0 as a function of time (Figs. 1C, 1D), which indicated that PKA stimulation led to rapid accumulation of FES-PKA at endomembranes. The effect was reversible with the treatment of 40 μM H89, a PKA inhibitor. A reverse order of drug treatment (H89, then FSK/IBMX) led to a similar result (Fig. S1). The balance of FES-PKA localization between the two subcellular locations (plasma membrane and endomembrane) remained largely the same across different expression levels (Fig. S2). As mentioned earlier, K-Ras4b undergoes multi-step modifications including farnesylation to its C-terminal region. To examine contribution of posttranslational modifications of FES-PKA for their membrane localization, we measured its localization in the presence of a farnesyltransferase inhibitor, lonafarnib. As a result, we observed a reduced number of cells with positive localization at the plasma membrane, along with concomitant increase in the number of cells where FES-PKA localized elsewhere such as the cytosol (Fig. S3). This suggests that the membrane-localized FES-PKA are likely farnesylated.

**Figure 1:**
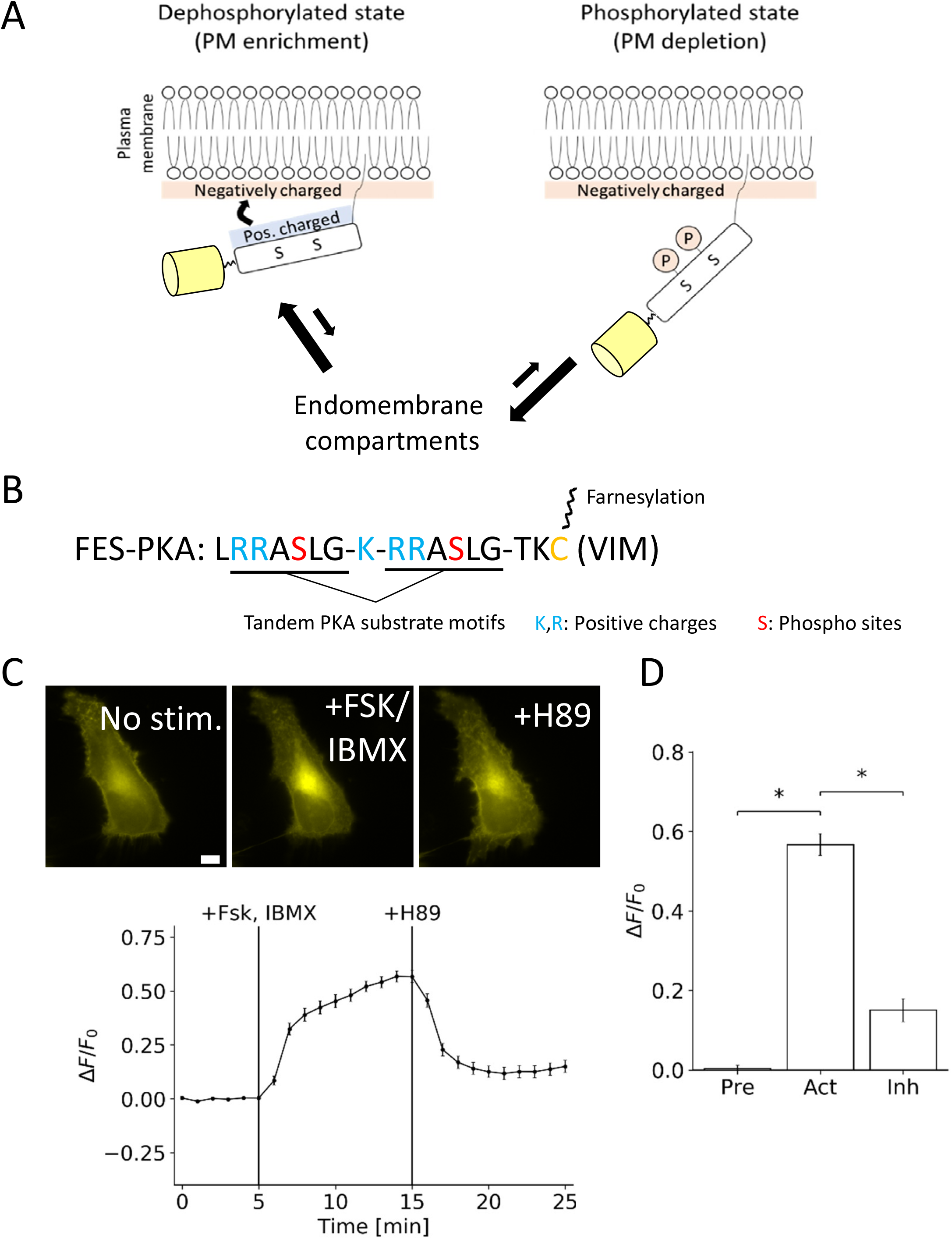
Basic Design and characterization of the Synthetic Farnesyl-Electrostatic Switch. **(A)** Plasma membrane targeting of farnesylated proteins relies on the presence of positively charged residues near the C-terminus. The farnesyl-electrostatic switch relies on the regulation of this sequence through phosphorylation. Yellow cylinders indicate a fluorescent protein for visualization. **(B)** Design of a farnesyl-electrostatic switch for PKA activity (FES-PKA) that consists of positive charges, two phosphate acceptors, and a CAAX tail which undergoes multi-step modifications including farnesylation. **(C)** Time-profile shows the quantification of the fluorescent signal in response to PKA activation and reversal with PKA inhibition. Representative micrographs of cells are shown on top in the following states: pre-treatment, post-activation, and post-inhibition. Fluorescence intensity of FES-PKA at the endomembranes at a given time point (*F*) was divided by that of time 0 (*F*0). The resulting value (*F*/*F*0) was then normalized to that of time 0, and used as an indicator of relative redistribution of FES-PKA in (C) and (D). All data points represent an average signal intensity collected from 30 cells over 3 independent experiments (10 cells each), and error bars represents standard error of mean. Scale bar represents 10 µm. **(D)** Bar chart represents the signal change pre-treatment (at 5 minutes), post-treatment with FSK+IBMX (at 15 minutes), and washout with H89 (at 25 minutes), respectively. ∗ represents statistical significance: p < 0.05.

We next explored the parameter space of this design to determine how different variables would affect the FES-PKA’s response and localization. The balance between the number of phosphorylation sites and the number and orientation of basic residues may determine the dynamic range of the FES-PKA’s response from PKA stimulation (Fig. 2A). To test this, we varied the number of the Kemptide sequence, as well as lysine residues between them. First, we generated a peptide with a single phosphorylation site (monovalent) and a peptide with three phosphorylation sites (trivalent). When co-transfected in cells, the monovalent peptide was largely localized in the intracellular membranes, perhaps due to an insufficient positive charge content/orientation. The trivalent peptide was observed at the plasma membranes along with other locations (likely nucleus). Quantification showed minor response from the monovalent peptide after PKA activation (Figs. 2B, 2C). We then modified the number of lysine residues in the linker region, and found that an increase in the number of lysines led to a concomitant decrease in the response from PKA activation (Figs. 2D, 2E). These results suggested that the best design for FES-PKA requires two Kemptide sequences aligned in tandem and connected with a single lysine residue.

**Figure 2:**
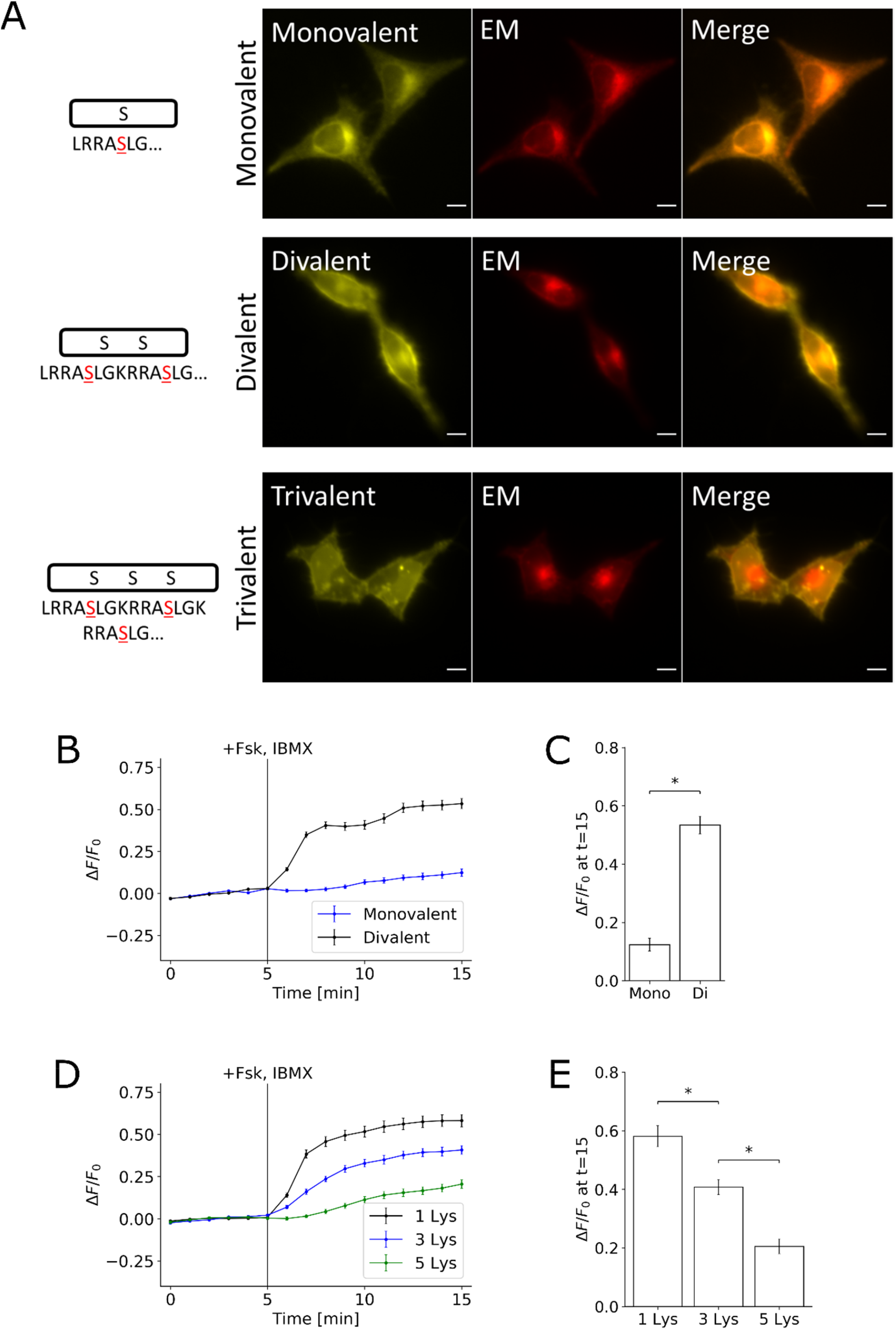
Effects of Substrate Valency and Positive Charge Content on FES-PKA Localization and Response. **(A)** Representative images are shown of cells co-transfected with peptides containing different numbers of substrates (monovalent, divalent, and trivalent) and an endomembrane (EM) marker. No stimulant was added. Divalent peptide is shown here as a reference and is the same amino sequence (FES-PKA) as shown in Fig. 1B. **(B)** Time-profile shows quantification of fluorescent signal of monovalent and divalent peptides in response to PKA activation. Trivalent peptide was not included in quantification due to localization defects. **(C)** Bar chart represents the signal change 10 minutes after PKA stimulation for the monovalent and divalent peptide. **(D)** Time-profile shows quantification of fluorescent signal of increasing the positive charge content of the peptide. **(E)** Bar chart represents the signal change 10 minutes after PKA stimulation for the three different lysine linkers. All data points in this figure represent an average signal intensity collected from 30 cells over 3 independent experiments (10 cells each), and error bars represents standard error of mean. ∗ represents statistical significance: p < 0.05. Scale bar represents 10 µm.

We next examined if the observed translocation of FES-PKA was due to phosphorylation and not due to perturbation of PKA activity by generating mutants which disrupted phosphorylation. Mutations of both serines in the optimized FES-PKA to alanines yielded little to no response (i.e., no translocation from PM to endomembranes upon PKA activation), while having both serines intact led to the most robust response (Figs. 3A, 3B). Having only one of the serines led to an intermediate response, suggesting that phosphorylation of each serine residue additively contributes to the switching mechanism.

**Figure 3:**
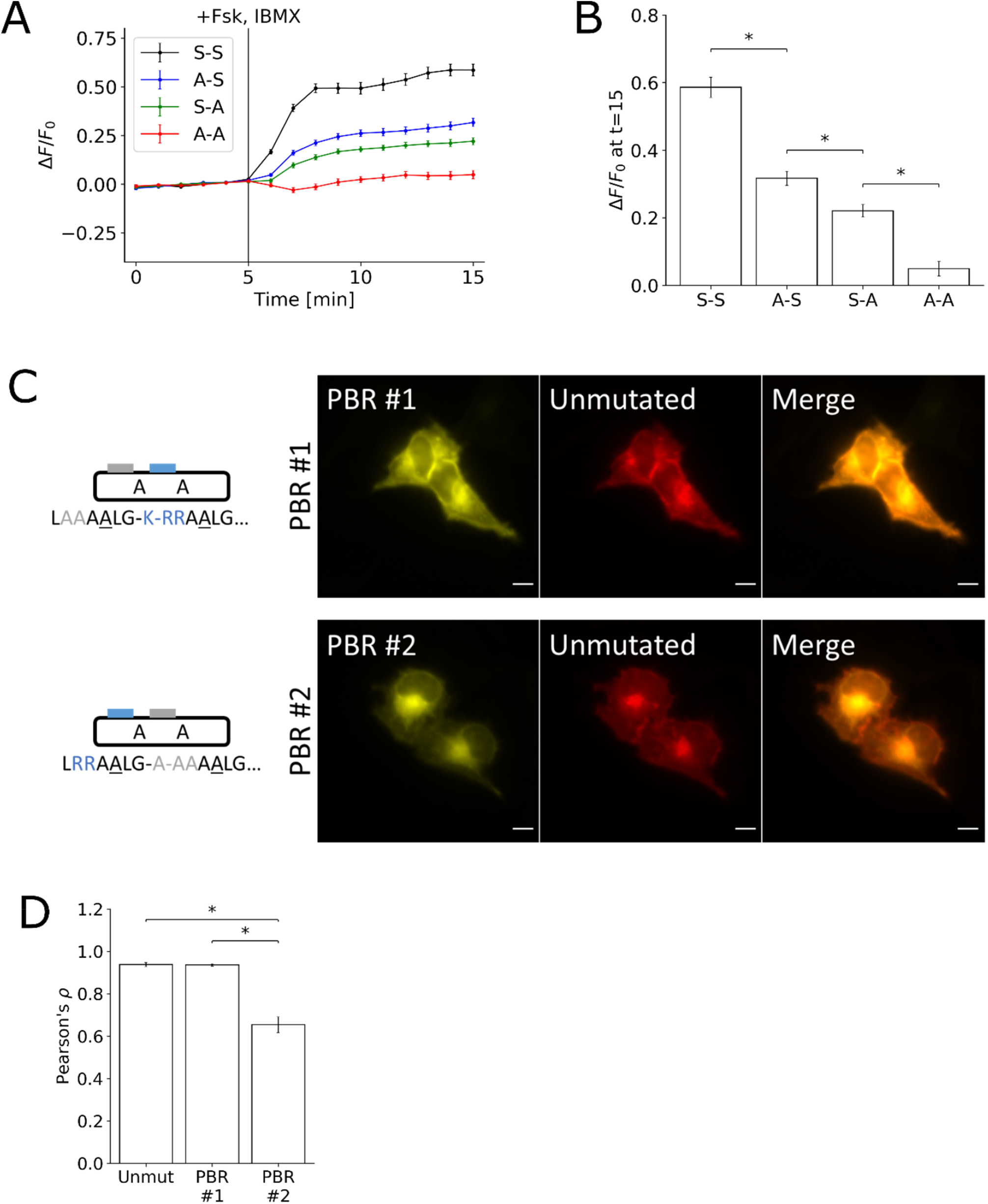
Role of Positively Charged Residues in Phosphorylation-Dependent Translocation. **(A)** Time-profile shows quantification of peptides with various permutations of non-phosphorylable FES-PKA. In the notation S-S, the first letter represents whether the first phosphorylation site is mutated, and the second serine represents whether the second phosphorylation site is mutated. **(B)** Bar chart represents the signal change for each of the mutations 10 minutes after PKA stimulation. **(C)** Micrograph represents cell images containing mutations disrupting the positively charged residues in the two clusters of basic residues, respectively. **(D)** Bar chart represents the average Pearson’s correlation calculated by comparing the three different peptides against their unmutated counterpart. All data points in Figs. 3A and 3B represent data collected from 30 cells over 3 independent experiments (10 cells each), and error bars represent standard error of mean. ∗ represents statistical significance: p < 0.05. Scale bar represents 10 µm.

In our design, we identified two clusters of basic residues, labeled as PBR #1 and PBR #2 (Fig. 3C). Based on the observation that the two serines are flanking PBR #2, we hypothesized that PBR #2 would have a more prominent role as a plasma membrane targeting sequence than PBR #1. We tested this hypothesis by mutating all the positively charged residues in PBR #1 and PBR #2 to alanines, respectively. As the arginines in P-2 and P-3 position are necessary for substrate recognition by PKA^21,24^, mutation of these residues to alanines disrupts the phosphorylability of the substrate. For this reason, the serines were mutated to alanines in this experiment. Cells were co-transfected with the mutated FES-PKA and a non-phosphorylable, non-mutated FES-PKA. Mutations in PBR #1 did not appear to disrupt the initial localization, while mutations in PBR #2 caused a substantial enrichment in the intracellular membranes (Fig. 3C). Pearson’s correlation was calculated by comparing the unmutated peptide against the three variant peptides (Fig. 3D).

Polybasic targeting sequences in farnesylated proteins including K-Ras4b can be disrupted with calcium (Ca^2+^) influx^15,25–28^. This is thought to occur through the binding of calmodulin, independent of phosphorylation. To test this potential effect of Ca^2+^, we elevated intracellular Ca^2+^ concentration by treating the cells with 1 μM ionomycin under a condition where the extracellular medium contained 1.8 mM Ca^2+^. The time-profile shows the translocation of FES-PKA (A-A), which is an unphosphorylable mutant (Fig. S4A). Both the S-S wild type and the A-A mutant, regardless of the presence of serines, responded similarly to the influx of Ca^2+^ (Fig. S4B). To further characterize the response to Ca^2+^, we used a Ca^2+^ chelator BAPTA while we monitored FES-PKA redistribution triggered by PKA activation. Despite the greatly suppressed Ca^2+^ with BAPTA, the extent and kinetics of FSK-IBMX-induced FES-PKA redistribution remained largely intact (Fig. S4C), suggesting that the observed response of FES-PKA is primarily driven by PKA activity, and contribution of Ca^2+^ is marginal.

Finally, we evaluated the validity of FES-PKA as a molecular sensor in relation to that of an existing PKA biosensor such as AKAR3EV^29^, a widely-used Förster resonance energy transfer (FRET) sensor. The fluorescence signal and the FRET signal were measured respectively in real time before and after stimulation with FSK/IBMX (Figs. 4A, 4B). As a result, FES-PKA indicated comparable dynamic range and kinetics to AKAR3EV (∼70% efficiency). Unlike FRET measurements that require additional optical setup and data computation, FES-PKA utilizes only a single fluorescence channel along with straightforward data analysis, which could find a wide, convenient use, as well as more advanced use such as multiplex imaging where activity of multiple biomolecules can be simultaneously visualized.

**Figure 4:**
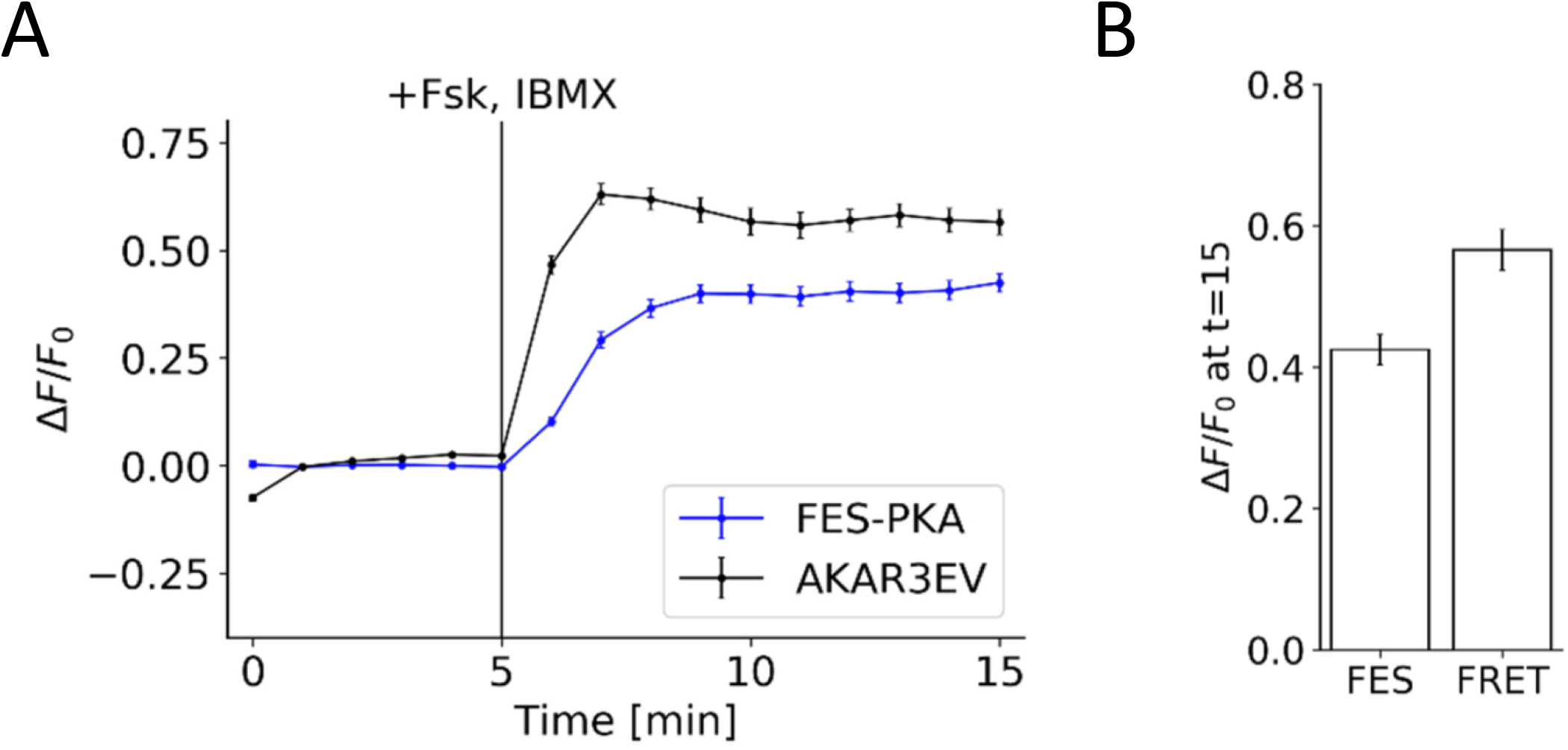
Comparison of FES-PKA to AKAR3EV. **(A)** Time-profile shows quantification of FES-PKA response in comparison to a PKA FRET sensor (AKAR3EV) in response to PKA activation. **(B)** Bar chart represents the signal change for the FES-PKA (FES) and AKAR3EV (FRET) 10 minutes after PKA activation. All data points represent an average signal intensity collected from 30 cells over 3 independent experiments (10 cells each), and error bars represents standard error of mean. ∗ represents statistical significance: p < 0.05. Scale bar represents 10 µm.

## Discussion

We engineered a PKA substrate peptide whose subcellular localization switches between the plasma membrane and the endomembrane depending on the cellular PKA activity. This molecular switch, termed FES-PKA, has further utility in future studies. For example, FES-PKA fused to a fluorescent protein can be used as a kinase activity sensor as demonstrated in Fig. 4. There have been reports on biosensors that change their intracellular localization in response to activation by an intended kinase, known as kinase-translocation reporters (KTRs). To our knowledge, all of these KTRs developed thus far shuttle within the soluble part of cells, namely between the nucleus and the cytosol^25-27^. This translocation is determined by the balance between nuclear export signal (NES) and nuclear localization signal (NLS) that can be modulated by a target kinase activity. Instead, the FES-PKA uses a unique switching mechanism whereby it travels within the membrane systems, potentially allowing for a multiplex usage with other KTRs without competing for cellular translocation machinery. It is also worth noting that the size of previous KTRs ranges from 60 to 400 amino acids which consist of multiple modules such as one for kinase binding, phosphorylation, NLS and NES. In contrast, FES-PKA is concise with only 20 amino acids due to employment of multi-tasking modules. As shown in Fig. S1, however, the FES-PKA responds not only to PKA, but also to Ca^2+^ as well as potentially other molecules, resulting in a tradeoff in target specificity.

Another application of FES-PKA would be to use it for a molecular actuator that can achieve conditional manipulation of biochemical activity. For example, one may conjugate FES-PKA to an enzyme for post-translational modification. Upon PKA activation, such a molecule could exert the enzymatic reaction due to increased proximity to its substrate as a result of altered subcellular localization. These enzymes include kinases and phosphatases for proteins and membrane lipids, as well as guanine exchange factors for small GTPases. As these molecular actuators could mediate cross-talk between two signal transduction pathways, synthetic operations such as generating a functional output in the face of a normally uncoupled input stimulus may become possible. In conclusion, due to the concise size and modular design principle, FES-PKA represents a potential for utilization in areas of biology and engineering.

## Materials and Methods

### Cell Culture and Transfection

HeLa (CCL-2, ATCC, Manassas, VA) cells were routinely passaged and cultured in DMEM (10-013-CV, Corning, Corning, NY) supplemented with 10% FBS (F2442, Sigma, St. Louis, MO) and maintained at 37°C in 5% CO_2_. For imaging experiments, cells were seeded at a 30-40% confluence on a #1 coverslip (48380-080, VWR, Radnor, PA) placed inside 6-wells (353046, Corning, Corning, NY). Cells were transfected the next day with FugeneHD (E2311, Promega, Madison, WI) at a ratio of 1:1 according to the manufacturer’s protocol, and 50 ng of DNA was added dropwise into each well. Cells were serum-starved with DMEM (17-205-CV, Corning, Corning, NY) without phenol red and supplemented with 2 mM L-glutamine (25-005-CL, Corning, Corning, NY) the morning of the following day. In the afternoon of the same day, cells were imaged by transferring the coverslip to an Attofluor cell chamber (A7816, Invitrogen, Carlsbad, CA).

### Image Acquisition

Images were acquired on an inverted microscope (IX81, Olympus, Tokyo, Japan) with a heated chamber that maintained conditions of 37°C and 5% CO_2_ (WELS, Tokai-Hit, Fujinomiya, Japan). Images were acquired for 15 minutes or 25 minutes, as noted, at 1-minute intervals. After 5 minutes, cells were treated with either 50 µM forskolin (F6886, Sigma, St. Louis, MO) and 100 µM IBMX (I5879, Sigma, St. Louis, MO) cocktail or 1 µM ionomycin (I-6800, LC Laboratories, Woburn, MA), as noted. Images were acquired with a CMOS camera (C11440, Hamamatsu Photonics, Hamamatsu, Japan) on a 60x oil objective (60x PlanApo N, Olympus, Tokyo, Japan). The microscope was controlled with MetaMorph (Molecular Devices, San Jose, CA) with motorized stage controller (MS-2000, ASI, Eugene, OR) and filter wheel controller (Lambda 10-3, Sutter Instrument, Novato, CA). The sample was illuminated with a LED light source (pE-300, CoolLED, Andover, UK). For FRET experiments, images were acquired by exciting with CFP and detecting in the YFP channel. All experiments shown in a given panel were performed together.

### Image Analysis

The signal at the intracellular membranes was quantified by co-expressing in cells an endomembrane marker that consisted of a prenylation sequence with no basic residues. ImageJ (NIH, Bethseda, MD) was used to track a 15×15 pixel (4.66 pixel = 1 μm) square region that contained the greatest average fluorescence intensity of the endomembrane marker. For fluorescence quantification, the background was subtracted, and the response of the signal was calculated by taking the ratio of signal with the average of the pre-treatment signal at the region of interest determined with the endomembrane marker. For FRET quantification, the background was subtracted, and the CFP-excited YFP emission signal was divided by the CFP-excited CFP emission signal. The reported response is this quantity normalized by the average of the pre-treatment signal for the whole cell.

### Colocalization Analysis

For colocalization analysis, the role of the two polybasic residue clusters was characterized by mutating the positively charged residues to alanines, respectively. Cells were co-transfected with the mutated peptide and an unmutated peptide, both unphosphorylable. The extent of the colocalization between the two peptides was analyzed by first calculating a 20×20 pixel median filtered image for each channel and respectively subtracting them from the original images. The two background-subtracted images were correlated using Pearson’s correlation. The average of these Pearson’s correlations from 30 cells is shown in Fig. 3D.

### Statistical Analysis

Paired Student’s t-test was used to test for statistical significance in Fig. 1D. Unequal variance t-test was used to test for statistical significance in Figs. 2C, 2E, 3B, 3D, and 4B. For each analysis, sample size was 30 cells collected over 3 independent experiments (10 cells each). All error bars represent mean ±SEM.

### DNA Cloning

DNA constructs were cloned by inserting annealed oligonucleotides into EYFP-C1 (Takara Bio, Kusatsu, Japan) and mCherry-C1 (Takara Bio, Kusatsu, Japan) at the SacII and BamHI site. The forward sequences of the inserted oligonucleotides with the restriction sites are as follows:

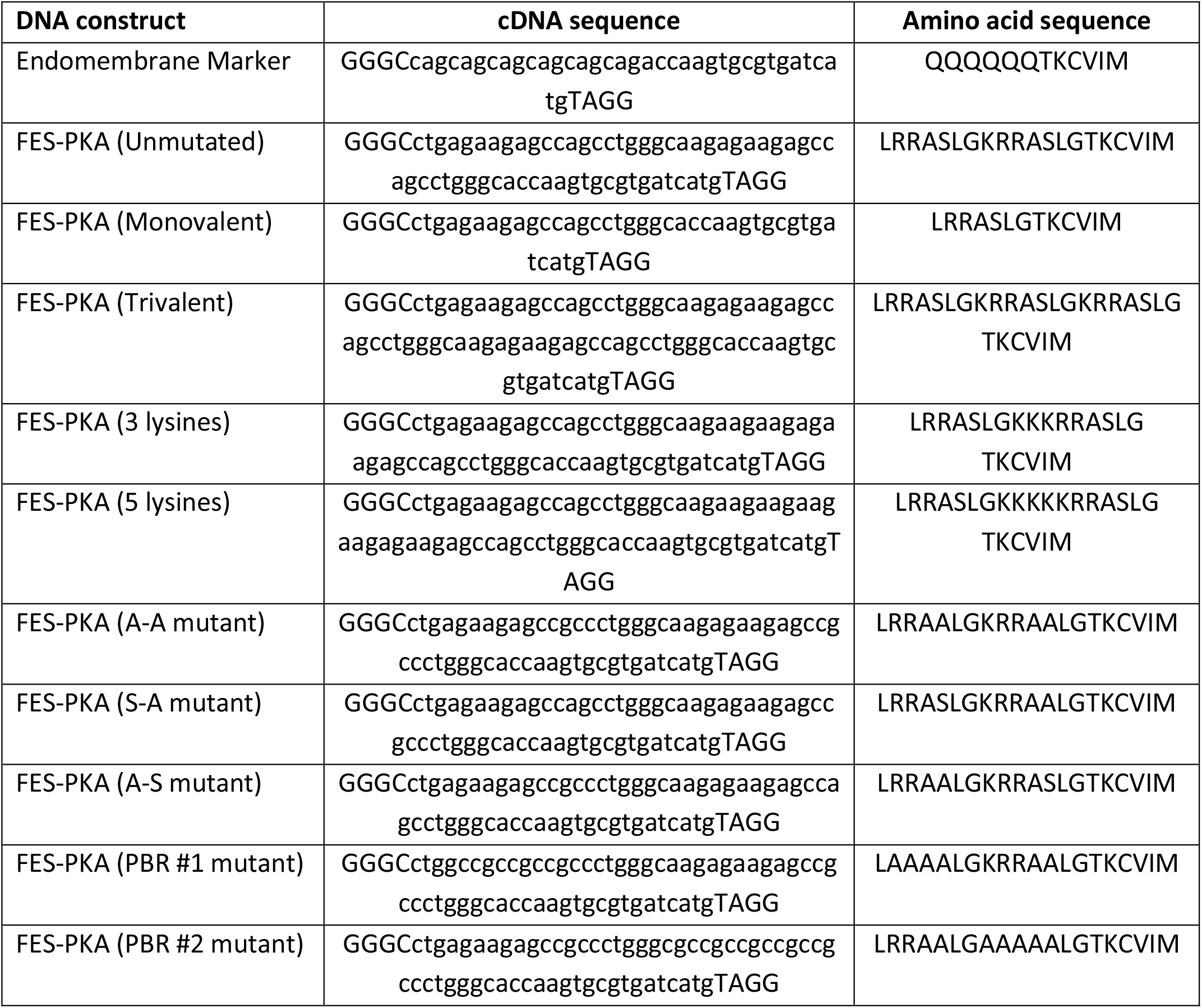

## Acknowledgments

We would like to thank Dr. Hideaki Matsubayashi for discussions that served as a starting point for the project and Dr. Abhijit Deb Roy for reading over the manuscript. We would also like to thank Dr. Kazuhiro Aoki for providing the AKAR3EV construct. This study was supported by National Institute of Health R01 GM123130 and DARPA HR0011-16-C-0139 (to T.I.).

## Author Contributions

T.I. and A.K. conceived and designed this study. A.K. and H.D.W. conducted experiments and analyses. A.K. and T.I. wrote the manuscript.

## Additional Information

The authors declare no competing interests.

## Notes

### Competing Interest Statement

The authors have declared no competing interest.

