## Supplementary Information for "Synthetic design of farnesyl-electrostatic peptides for development of a protein kinase A membrane translocation switch"

### Contents:

- Legend for Supplementary Movie 1
- Supplementary Figure 1-4 and their figure legends

Legend for Supplementary Movie 1

HeLa cells were transfected with FES-PKA as a YFP fusion whose fluorescence was measured every minute while cells were stimulated with FSK/IBMX for 10 min, and then with H89 for 10 min. The cell shown corresponds to the one in Fig. 1C.

Figure S1

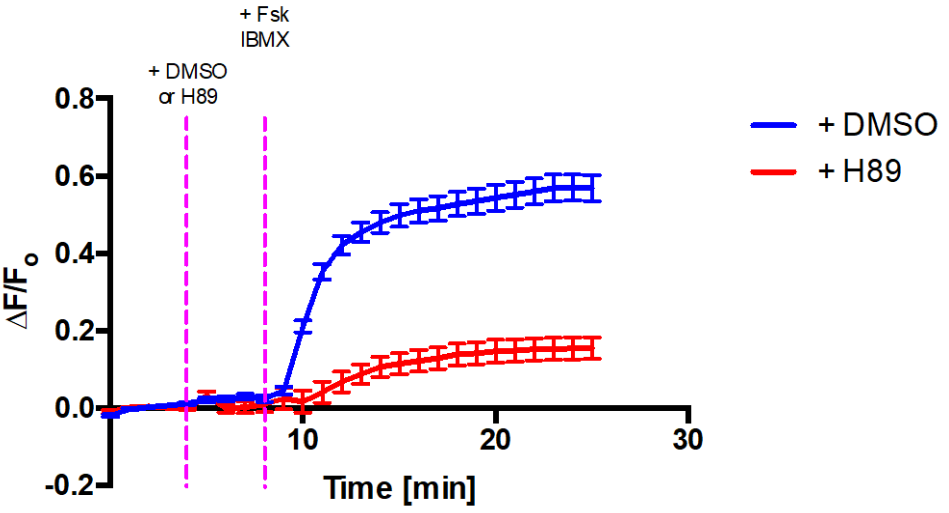


**<u>Figure S1:</u> H89 treatment prior to PKA activation by FSK/IBMX.** Graph shows quantification of time-lapse imaging of HeLa cells co-expressing FES-PKA and EM marker. 4 mins after image acquisition, 40 uM H89 or 0.1% DMSO was added, followed by the of 50 μM Fsk / 100 uM IBMX at 9 mins. ($P < 0.0001$, N = 3 experiments/condition, 62 cells/DMSO, 35 cells/H89)

Figure S2

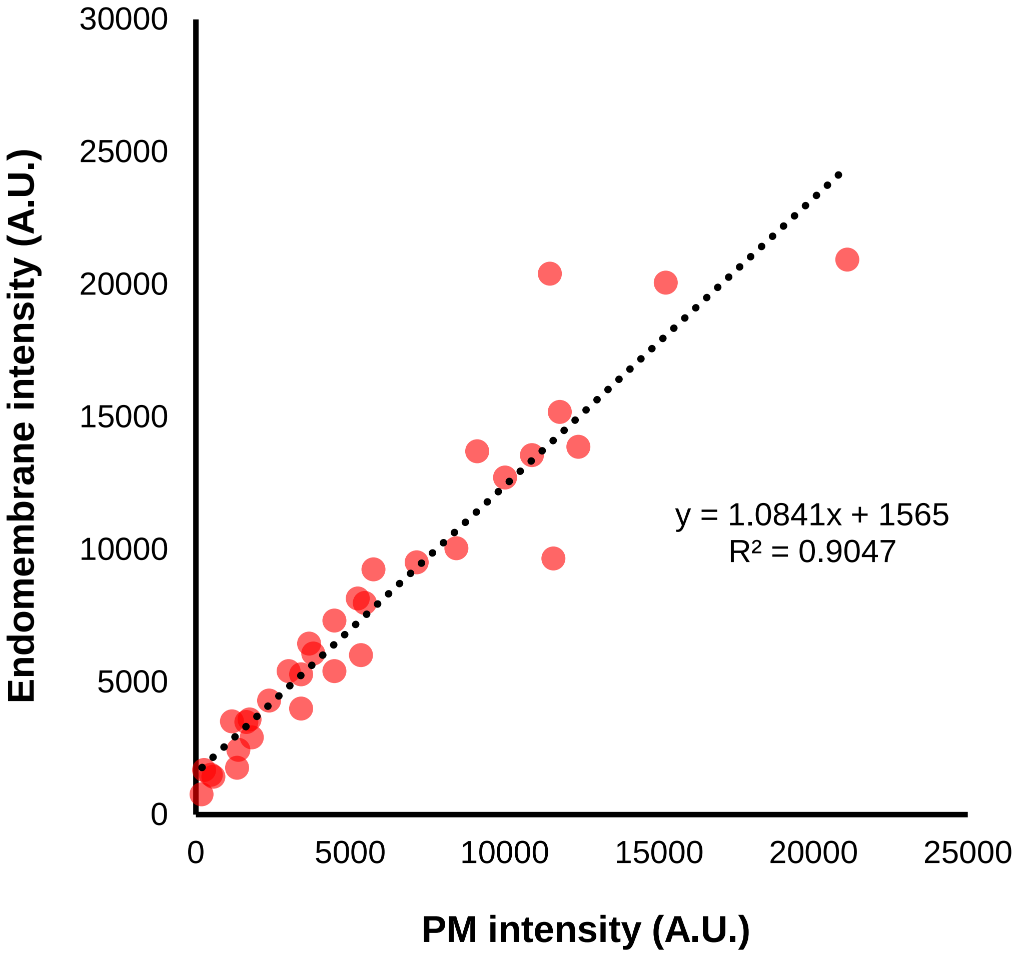


**<u>Figure S2:</u> Effects of expression level on PM and endomembrane intensity.** FES-PKA intensity at PM and endomembrane are determined in HeLa cells also co-expressing EM marker and PM marker (Lyn-CFP). PM localization was taken by line-scan analysis. EM localization was determined at a region (20 by 20 pixel) of maximum EM intensity. (N = 3 experiments, 33 cells)

Figure S3

A

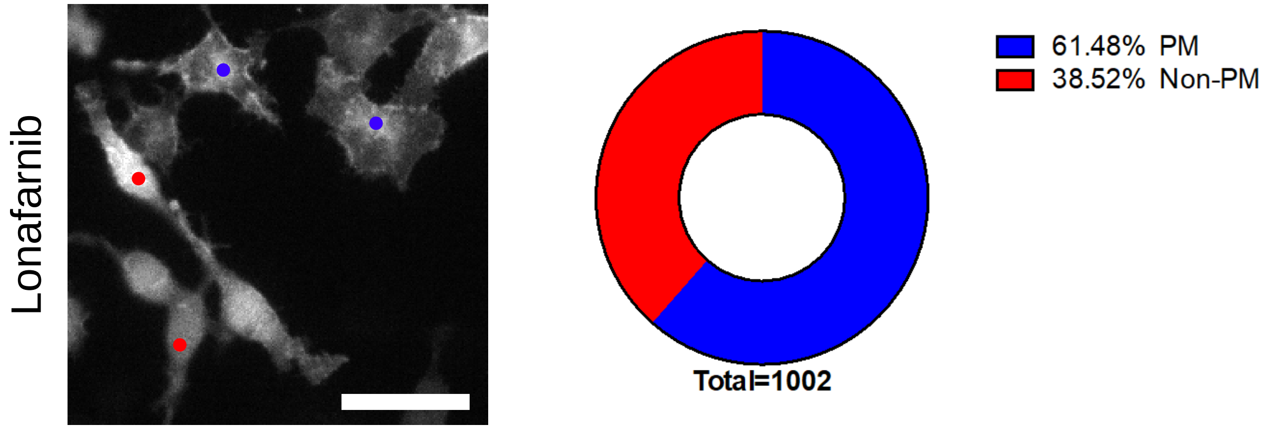


B

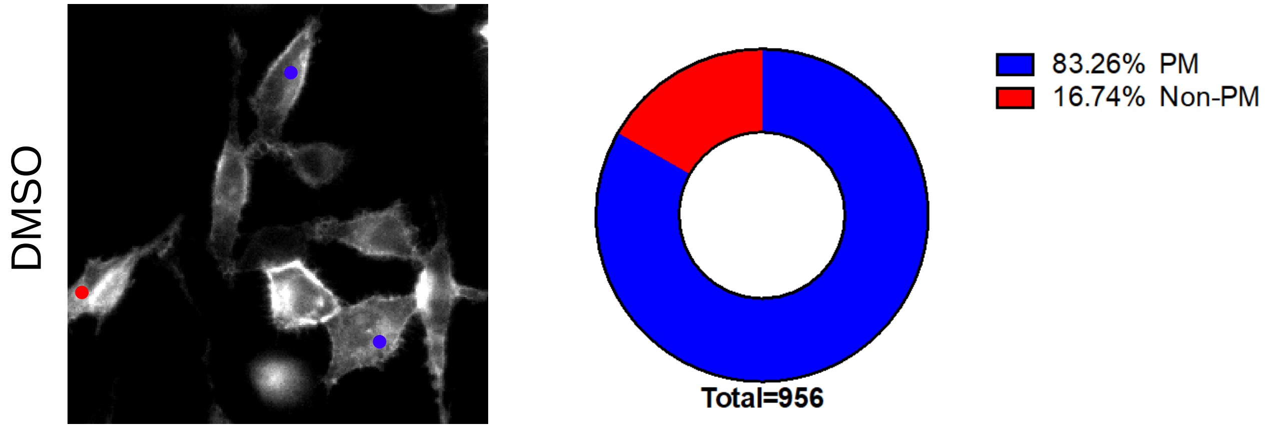


C

| | Non-PM | Total | Risk | Risk Ratio | 95 % CI |
|---|---|---|---|---|---|
| **Lonafarnib** | 386 | 1002 | 0.385 | 2.30 | 2.55 – 2.07 |
| **DMSO** | 160 | 956 | 0.167 | | |

**<u>Figure S3:</u> Effects of farnesyltransferase inhibition on FES-PKA localization.** HeLa cells co-expressing FES-PKA and EM marker treated overnight with (A) 2 μM Lonafarnib or (B) 0.1% DMSO. Localization to the PM (blue) or Non-PM (red) as classified by eye. Scale bar, 50 μm. (C) Risk for non-PM localization for each condition (Lonafarnib or DMSO) is non-PM/total. Risk ratio is Lonafarnib risk divided by DMSO risk. 95% confidence interval (CI) is based on the equation 95% CI = exp (ln RR ± SQRT (1/a + 1/b + 1/c + 1/d), where a = number of cells showing non-PM FES-PKA localization (lonafarnib), b = total number of cells analyzed (lonafarnib), c = number of cells showing non-PM FES-PKA localization (DMSO), d = total number of cells analyzed (DMSO). Because 95% CI does not include the null value (RR = 1), lonafarnib significantly increased non-PM localization of FES-PKA ($P < 0.05$). (N = 3 experiments / condition.)

Figure S4

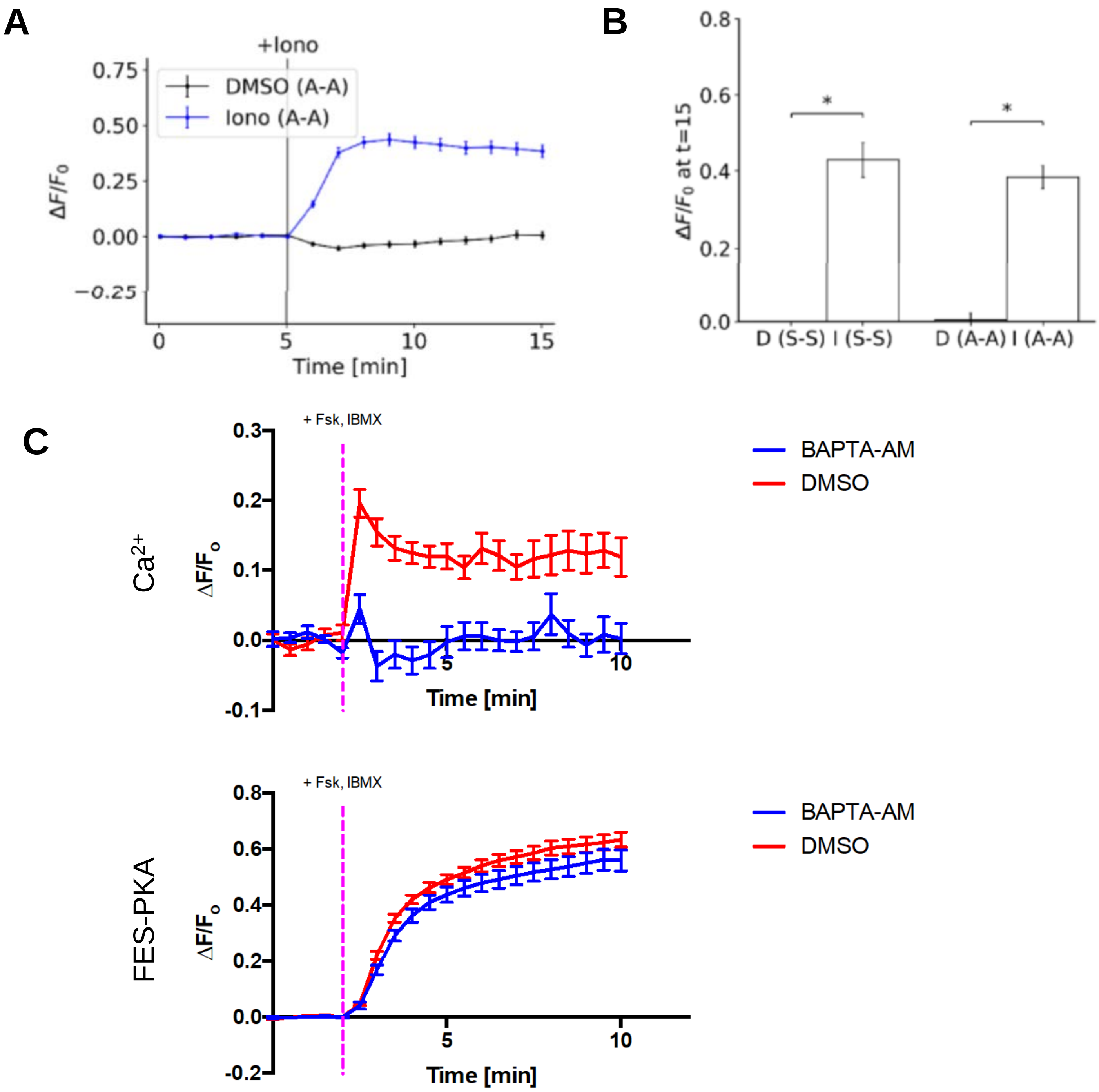


**<u>Figure S4:</u> Effects of intracellular calcium on FES-PKA translocation.** (A) Time-profile shows quantification of a non-phosphorylable FES-PKA in response to $Ca^{2+}$ influx. (B) Bar chart represents the signal change of a non-phosphorylable FES-PKA (A-A) and phosphorylable FES-PKA (S-S), respectively, after 10 minutes of treatment with DMSO (D) and ionomycin (I), respectively. (C) Cells co-expressing FES-PKA and R-GECO calcium indicator are pre-loaded with 0.08% Pluronic F-127 with 30 μM BAPTA-AM intracellular calcium chelator or 0.4% DMSO. Top panel shows calcium response (R-GECO intensity) to 50 μM Fsk / 100 uM IBMX added at 2 mins after image acquisition ($P < 0.0001$). Bottom panel indicates FES-PKA translocation response to Fsk/IBMX added 2 mins after image acquisition ($P = 0.548$). (N = 3 experiments/condition, 26 cells/BAPTA-AM, 36 cells/DMSO)
